# CeTF: an R package to Coexpression for Transcription Factors using Regulatory Impact Factors (RIF) and Partial Correlation and Information (PCIT) analysis

**DOI:** 10.1101/2020.03.30.015784

**Authors:** Carlos Alberto Oliveira de Biagi, Ricardo Perecin Nociti, Breno Osvaldo Funicheli, Patrícia de Cássia Ruy, João Paulo Bianchi Ximenez, Wilson Araújo Silva

## Abstract

**Summary:** Finding meaningful gene-gene associations and the main Transcription Factors (TFs) in co-expression networks is one of the most important challenges in gene expression data mining. CeTF is an R package that integrates the Partial Correlation with Information Theory (PCIT) and Regulatory Impact Factors (RIF) algorithms applied to gene expression data from microarray, RNA-seq, or single-cell RNA-seq platforms. This approach allows identifying the transcription factors most likely to regulate a given network in different biological systems — for example, regulation of gene pathways in tumor stromal cells and tumor cells of the same tumor. This pipeline can be easily integrated into the high-throughput analysis.

**Availability:** CeTF is available as R package in Bioconductor (https://bioconductor.org/packages/CeTF), GitHub (https://github.com/cbiagii/CeTF) and as docker image (https://hub.docker.com/r/biagii/cetf). More information on how to use the package can be found in the Supplemental File 1.

## 1. Introduction

Gene expression data analysis has become crucial to biological sciences, one of the most interesting forms of analyzing this type of data is the gene-to-gene network interaction analysis, aiming to highlight which gene interactions are the most relevant to the study. Despite the plethora of tools, new methods are needed to evaluate all possible interactions and their significance (Yu et al., 2013). In addition to identifying pairwise gene-to-gene interaction, finding the transcription factors (TFs) is of great interest, mainly because they can play an essential regulatory role (Farnham, 2009). Furthermore, integrating network generation with the identification of main TFs brings a deciding view of the data. In this article, we provide an R package that enables the performing of the Regulatory Impact Factors (RIF) and Partial Correlation with Information Theory (PCIT) analysis separately, or by applying the full pipeline. This package will be useful for creating a network from gene expression data identifying the most significant pairwise interactions and main TFs.

## 2. Implementation and main functions

CeTF is an implementation in R for PCIT (Reverter and Chan, 2008) and RIF (Reverter et al., 2010) algorithms, which initially were made in FORTRAN language. From these two algorithms, it was possible to integrate them in order to increase performance and results. Input data may come from microarray, RNA-seq, or single-cell RNA-seq. As seen earlier, the input data can be read counts or expression (TPM, FPKM, normalized values, and others). The main pipeline (Fig. 1) consists of the following steps.

**Fig. 1:**
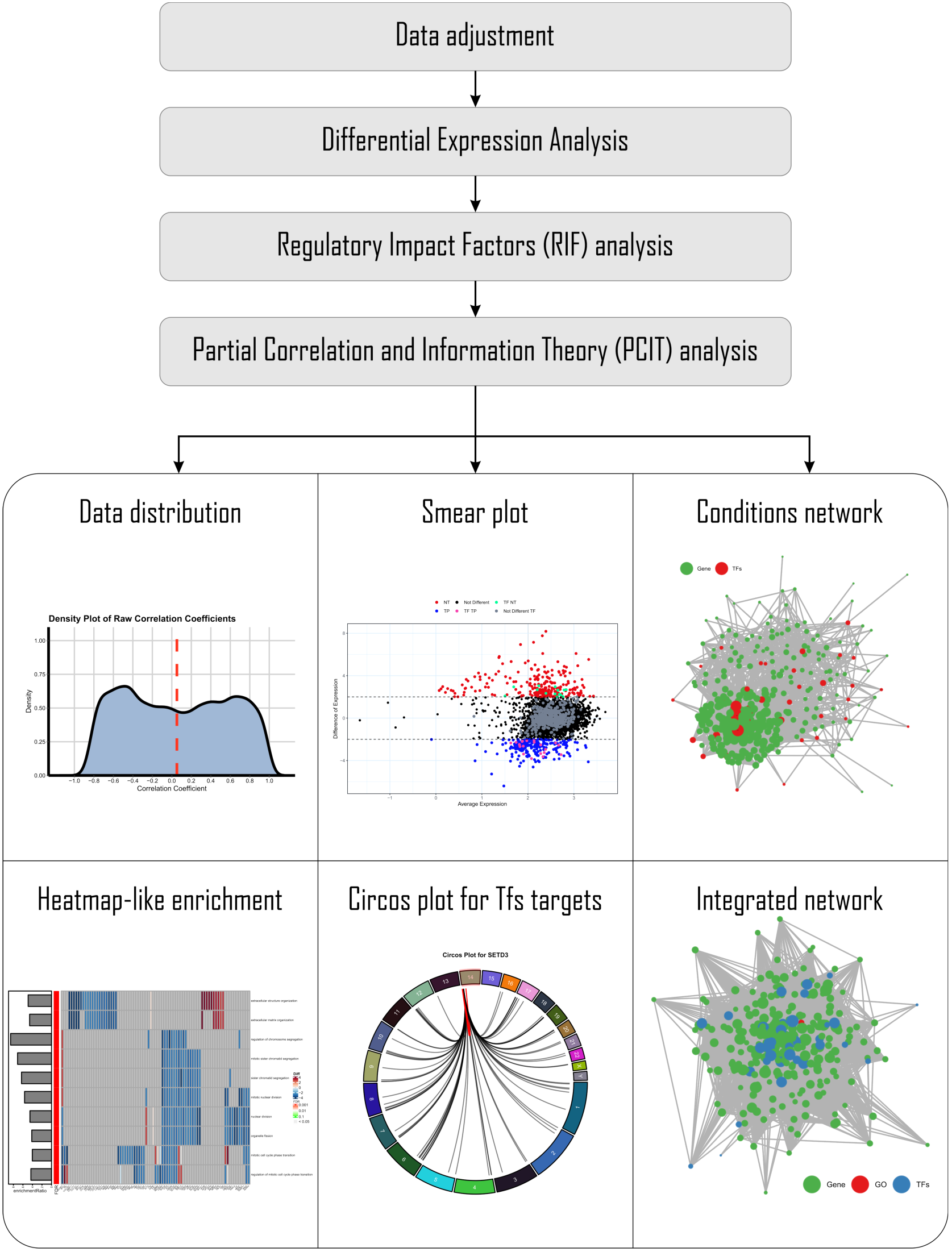
Schematic of a CeTF workflow. The main functions of CeTF and their logical order are illustrated. Grey boxes signify the mandatory steps for the main analysis. The white box signifies the optional outputs from CeTF package, i.e. data distribution, smear plot, conditions network, heatmap-like enrichment, circus plot for TFs targets, integrated network, etc.

### Step 1: Data adjustment

If the input data is a count table, data will be converted to TPM by each column (x) as follows

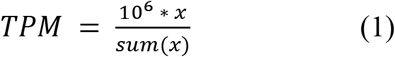

The mean for TPM values different than zero, and the mean values for each gene are used as a threshold to filter the genes. Genes with values above half of the previous averages will be considered for subsequent analyses. Then, the TPM data is normalized using:

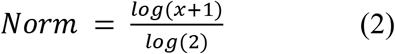

If the input already has normalized expression data (TPM, FPKM, etc), the only step will be the same filter for genes that consider half of the means.

### Step 2: Differential Expression analysis

For differential analysis of gene expression, there are two options, the *Reverter* method (Reverter *et al.*, 2006) and DESeq2 (Love *et al.*, 2014). In both methods, two conditions are required (i.e., control *vs.* tumor samples). In the *Reverter* method, the mean between samples of each condition for each gene is calculated. Then, subtraction is made between the mean of one condition concerning the other conditions. The variance of the subtraction is performed, then is calculated the difference of expression using the following formula, where *s* is the result of subtraction and *var* is the variance:

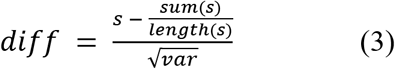

The DESeq2 method applies the differential expression analysis based on the negative binomial distribution. Although both methods can be used on count data, it is strongly recommended to use only the *Reverter* method on expression input data.

### Step 3: Regulatory Impact Factors (RIF) analysis

The RIF algorithm is well described in the original paper (Reverter *et al.*, 2010). This step aims to identify critical Transcription Factors calculating for each condition the co-expression correlation between the TFs and the Differentially Expressed (DE) genes (from Step 2). The result is RIF1 and RIF2 metrics that allow the identification of critical TFs. The RIF1 metric classifies the TFs as most differentially co-expressed with the highly abundant and highly DE genes, and the RIF2 metric classifies the TF with the most altered ability to act as predictors of the abundance of DE genes. A main TF is defined if:

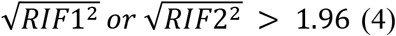

### Step 4: Partial Correlation and Information Theory (PCIT) analysis

The PCIT algorithm is also well described in the original paper from Reverter and Chan (Reverter and Chan, 2008). Moreover, it has been used for the reconstruction of Gene Co-expression Networks (GCN). The GCN combines the concept of Partial Correlation coefficient with Information Theory to identify significant gene-to-gene associations defining edges in the reconstruction of the network. At this stage, the paired correlation of three genes is performed at the same time, thus making the inference of co-expressed genes. This approach is more sensitive than other methods and allows the detection of functionally validated gene-gene interactions. First, is calculated for every trio of genes x, y, and z the partial correlation coefficients:

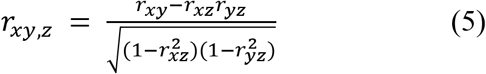

And similarly, for *r*_*xz,y*_ and *r*_*yz,x*_. After that, for each trio of genes is calculated the tolerance level (*ε*) to be used as a threshold for capturing significant associations. The average ratio of partial to direct correlation is computed as follows:

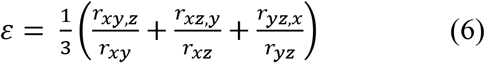

The association between the genes *x* and *y* is discarded if:

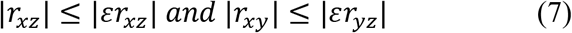

Otherwise, the association is defined as significant, and the interaction between the genes *x* and *y* is used in the reconstruction of the GCN.

The final output includes the network with gene-gene and gene-TF interactions for both conditions, besides generating the main TFs identified in the network.

## 3. Additional functionalities

The CeTF package also has some additional features that includes plots to visualize the data distribution, the distribution of differentially expressed genes/TFs that shows the average expression (in log2) by the difference of expression, and the network for both conditions. It is also possible to perform the grouping of ontologies (Carbon *et al.*, 2019) without statistical inference and functional enrichment for several databases with statistical inference of any organism. Finally, the network that integrates the genes, TFs and pathways is generated.

## 3. Conclusions

CeTF is a tool that assists the identification of meaningful gene-gene associations and the main TFs in co-expression networks. It offers functions for a complete and customizable workflow from count or expression data to networks and visualizations in the form of a freely available R package. We expect that CeTF will be widely used by the genomics and transcriptomics community and scientists that work with high-throughput data that aims to understand how main TFs are working in a co-expression network and what are the pathways involved in this context.

