## Supplemental File 1 for "CeTF: an R package to Coexpression for Transcription Factors using Regulatory Impact Factors (RIF) and Partial Correlation and Information (PCIT) analysis"

#### Introduction

This vignette provides the necessary instructions for performing the Partial Correlation coefficient with Information Theory (PCIT) (Reverter and Chan 2008) and Regulatory Impact Factors (RIF) (Reverter et al. 2010) algorithm.

The PCIT algorithm identifies meaningful correlations to define edges in a weighted network and can be applied to any correlation-based network including but not limited to gene co-expression networks, while the RIF algorithm identify critical transcript factors (TF) from gene expression data.

These two algorithms when combined provide a very relevant layer of information for gene expression studies (Microarray, RNA-seq and single-cell RNA-seq data).

##### Regulatory Information Factors (RIF)

A gene expression data from microarray, RNA-seq or single-cell RNA-seq spanning two biological conditions of interest (e.g. normal/tumor, healthy/disease, malignant/nonmalignant) is subjected to standard normalization techniques and significance analysis to identify the target genes whose expression is differentially expressed (DE) between the two conditions. Then, the regulators (e.g. Transcript Factors genes) are identified in the data. The TF genes can be obtained from the literature (Wang and Nishida 2015)(Vaquerizas et al. 2009). Next, the co-expression correlation between each TF and the DE genes is computed for each of the two conditions. This allows for the computation of the differential wiring (DW) from the difference in co-expression correlation existing between a TF and a DE genes in the two conditions. As a result, RIF analysis assigns an extreme score to those TF that are consistently most differentially co-expressed with the highly abundant and highly DE genes (case of RIF1 score), and to those TF with the most altered ability to act as predictors of the abundance of DE genes (case of RIF2 score). A given TF may not show a change in expression profile between the two conditions to score highly by RIF as long as it shows a big change in co-expression with the DE genes. To this particular, the profile of the TF gene (triangle, solid line) is identical in both conditions (slightly downwards). Instead, the DE gene (circle, dashed line) is clearly over-expressed in condition B. Importantly, the expression of the TF and the DE gene shows a strong positive correlation in condition A, and a strong negative correlation in condition B.

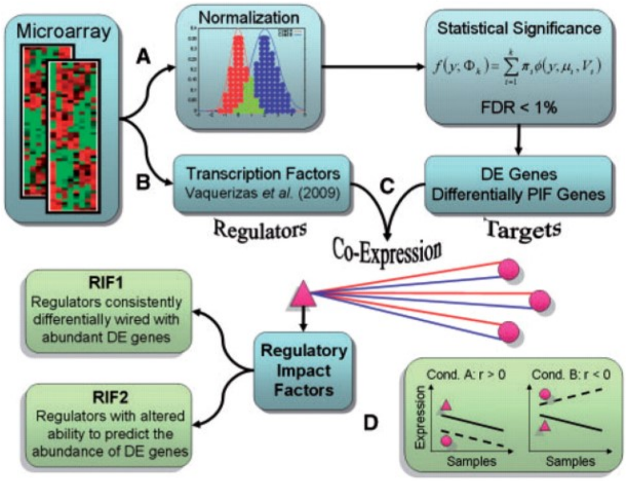


#### Partial Correlation with Information Theory (PCIT)

The proposed PCIT algorithm contains two distinct steps as follows:

**Step 1 - Partial correlations** For every trio of genes in x, y and z, the three first-order partial correlation coefficients are computed by:

$$r_{xy.z} = \frac{r_{xy} - r_{xz}r_{yz}}{\sqrt{(1 - r_{xz}^2)(1 - r_{yz}^2)}}$$

, and similarly for $r_{xz.y}$ and $r_{yz.x}$.

The partial correlation coefficient between $x$ and $y$ given $z$ (here denoted by $r_{xy.z}$) indicates the strength of the linear relationship between $x$ and $y$ that is independent of (uncorrelated with) $z$. Calculating the ordinary (or unconditional or zero-order) correlation coefficient and comparing it with the partial correlation, we might see that the association between the two variables has been sharply reduced after eliminating the effect of the third variable.

**Step 2 - Information theory** We invoke the Data Processing Inequality (DPI) theorem of Information Theory which states that 'no clever manipulation of the data can improve the inference that can be made from the data' (Cover and Thomas 2012). For every trio of genes, and in order to obtain the tolerance level ($\varepsilon$) to be used as the local threshold for capturing significant associations, the average ratio of partial to direct correlation is computed as follows:

$$\varepsilon = (\frac{r_{xy.z}}{r_{xy}} + \frac{r_{xz.y}}{r_{xz}} + \frac{r_{yz.x}}{r_{yz}})$$

In the context of our network reconstruction, a connection between genes $x$ and $y$ is discarded if:

$$|r_{xy}| \leq |\varepsilon r_{xz}| and |r_{xy}| \leq |\varepsilon r_{yz}|$$

Otherwise, the association is defined as significant, and a connection between the pair of genes is established in the reconstruction of the GCN. To ascertain the significance of the association between genes $x$ and $y$, the above mentioned Steps 1 and 2 are repeated for each of the remaining $n{-}2$ genes (denoted here by $z$).

#### Installation

To install, just type:

```
BiocManager::install("cbiagii/CeTF")
```

for Linux users is necessary to install **libcurl4-openssl-dev**, **libxml2-dev** and **libssl-dev** dependencies.

#### Workflow

There are many ways to perform the analysis. The following sections will be splited by steps, and finishing with the complete analysis with visualization. We will use the **airway** (Himes et al. 2014) dataset in the following sections. This dataset provides a RNA-seq count data from four human ASM cell lines that were treated with dexamenthasone - a potent synthetic glucocorticoid. Briefly, this dataset has 4 samples untreated and other 4 samples with the treatment.

##### PCIT

The first option is to perform the PCIT analysis. The output will be a list with 3 elements. The first one contains a dataframe with the pairwise correlation between genes (corr1) and the significant pairwise correlation (corr2 $\neq 0$). The second element of the list stores the adjacency matrix with all correlation. And the last element contains the adjacency matrix with only the significant values:

```
# Loading packages
library(CeTF)
library(airway)
library(kableExtra)
library(knitr)

# Loading airway data
data("airway")

# Creating a variable with annotation data
anno <- as.data.frame(colData(airway))
anno <- anno[order(anno$dex, decreasing = TRUE), ]
anno <- data.frame(cond = anno$dex,
                   row.names = rownames(anno))

# Creating a variable with count data
counts <- assay(airway)

# Sorting count data samples by conditions (untrt and trt)
counts <- counts[, rownames(anno)]
colnames(counts) <- paste0(colnames(counts), c(rep("_untrt", 4), rep("_trt", 4)))

# Differential Expression analysis to use only informative genes
DEGenes <- expDiff(exp = counts,
                   anno = anno,
                   conditions = c('untrt', 'trt'),
                   lfc = 4.5,
                   padj = 0.05,
                   diffMethod = "Reverter")

# Selecting only DE genes from counts data
counts <- counts[rownames(DEGenes$DE_unique), ]
```

```r
# Converting count data to TPM
tpm <- apply(counts, 2, function(x) {
            (1e+06 * x)/sum(x)
        })

# Count normalization
PCIT_input <- normExp(tpm)

# PCIT input for untrt
PCIT_input_untrt <- PCIT_input[,grep("_untrt", colnames(PCIT_input))]

# PCIT input for trt
PCIT_input_trt <- PCIT_input[,grep("_trt", colnames(PCIT_input))]

# Performing PCIT analysis for untrt condition
PCIT_out_untrt <- PCIT(PCIT_input_untrt, tolType = "mean")

# Performing PCIT analysis for trt condition
PCIT_out_trt <- PCIT(PCIT_input_trt, tolType = "mean")

# Printing first 10 rows for untrt condition
kable(PCIT_out_untrt$tab[1:10, ]) %>%
  kable_styling(bootstrap_options = "striped", full_width = FALSE)
```

| gene1 | gene2 | corr1 | corr2 |
|---|---|---:|---:|
| ENSG00000011465 | ENSG00000026025 | 0.01160 | 0.00000 |
| ENSG00000011465 | ENSG00000035403 | 0.61286 | 0.00000 |
| ENSG00000011465 | ENSG00000049323 | -0.71685 | 0.00000 |
| ENSG00000011465 | ENSG00000067225 | -0.99935 | -0.99935 |
| ENSG00000011465 | ENSG00000071127 | -0.56817 | 0.00000 |
| ENSG00000011465 | ENSG00000071242 | -0.89063 | 0.00000 |
| ENSG00000011465 | ENSG00000075624 | 0.09820 | 0.00000 |
| ENSG00000011465 | ENSG00000077942 | -0.80938 | 0.00000 |
| ENSG00000011465 | ENSG00000080824 | -0.72639 | 0.00000 |
| ENSG00000011465 | ENSG00000087086 | -0.91357 | 0.00000 |

```r
# Printing first 10 rows for trt condition
kable(PCIT_out_trt$tab[1:10, ]) %>%
  kable_styling(bootstrap_options = "striped", full_width = FALSE)
```

| gene1 | gene2 | corr1 | corr2 |
|---|---|---:|---:|
| ENSG00000011465 | ENSG00000026025 | 0.63800 | 0.00000 |
| ENSG00000011465 | ENSG00000035403 | 0.96857 | 0.96857 |
| ENSG00000011465 | ENSG00000049323 | 0.14794 | 0.00000 |
| ENSG00000011465 | ENSG00000067225 | -0.99687 | -0.99687 |
| ENSG00000011465 | ENSG00000071127 | -0.92734 | -0.92734 |
| ENSG00000011465 | ENSG00000071242 | -0.75914 | 0.00000 |
| ENSG00000011465 | ENSG00000075624 | -0.70489 | 0.00000 |
| ENSG00000011465 | ENSG00000077942 | -0.29712 | 0.00000 |
| ENSG00000011465 | ENSG00000080824 | -0.93148 | 0.00000 |
| ENSG00000011465 | ENSG00000087086 | -0.94620 | 0.00000 |

```r
# Adjacency matrix: PCIT_out_untrt$adj_raw or PCIT_out_trt$adj_raw
# Adjacency matrix only with the significant values: PCIT_out_untrt$adj_sig or PCIT_out_trt$adj_sig
```

#### Histogram of connectivity distribution

After performing the PCIT analysis, it is possible to verify the histogram distribution of the clustering coefficient of the adjacency matrix with the significant values:

```r
# Example for trt condition
histPlot(PCIT_out_trt$adj_sig)
```

```
## Warning: Use of `df1[["clustcoef"]]` is discouraged. Use `.data[["clustcoef"]]`
## instead.

## Warning: Use of `df2[["clustcoef"]]` is discouraged. Use `.data[["clustcoef"]]`
## instead.
```

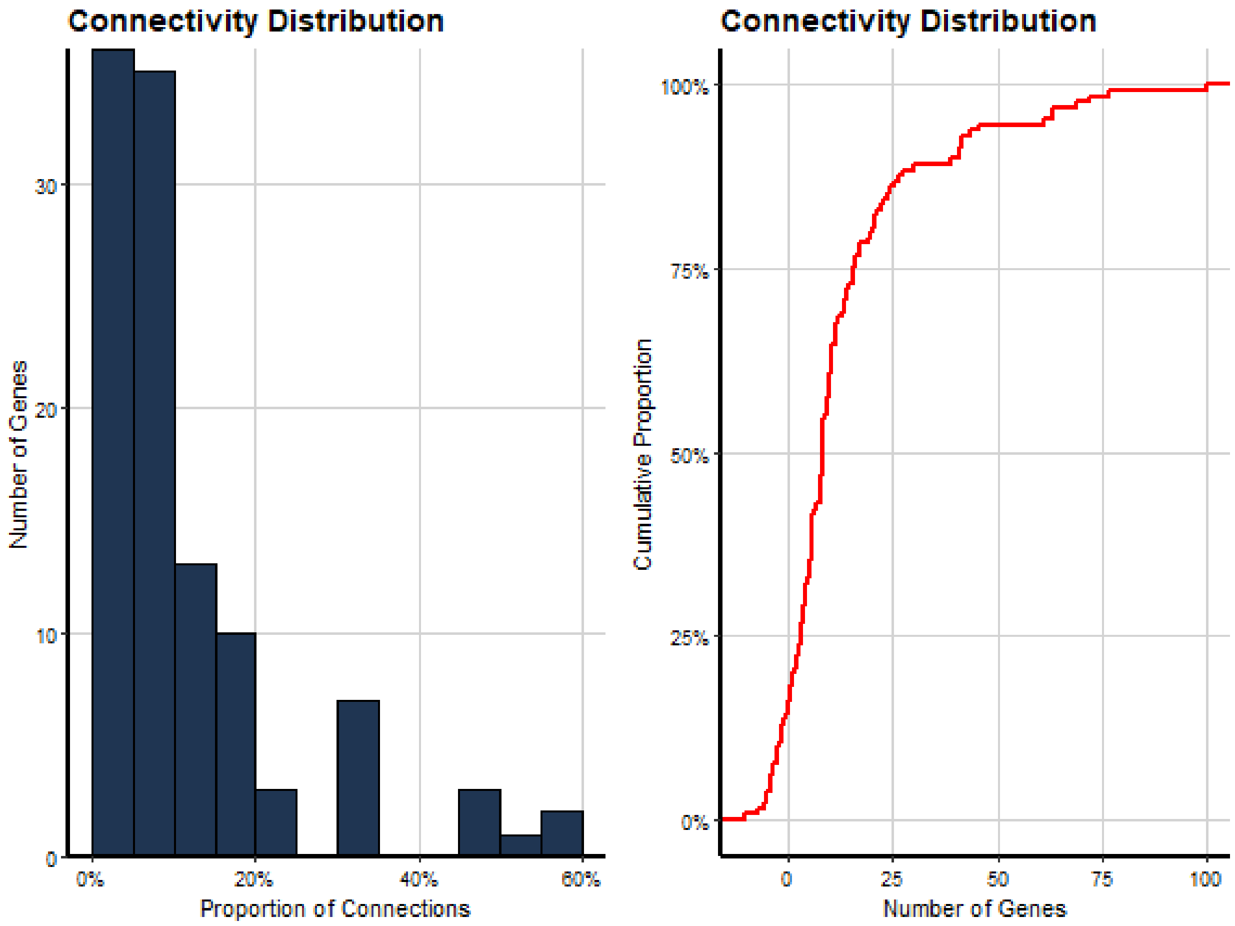


#### Density Plot of raw correlation and significant PCIT

It's possible to generate the density plot with the significance values of correlation. We'll use the raw adjacency matrix and the adjacency matrix with significant values. It is necessary to define a cutoff of the correlation module (values between -1 and 1) that will be considered as significant:

```r
# Example for trt condition
densityPlot(mat1 = PCIT_out_trt$adj_raw,
            mat2 = PCIT_out_trt$adj_sig,
            threshold = 0.5)
```

```
## Warning: Use of `df1[["corr"]]` is discouraged. Use `.data[["corr"]]` instead.
```

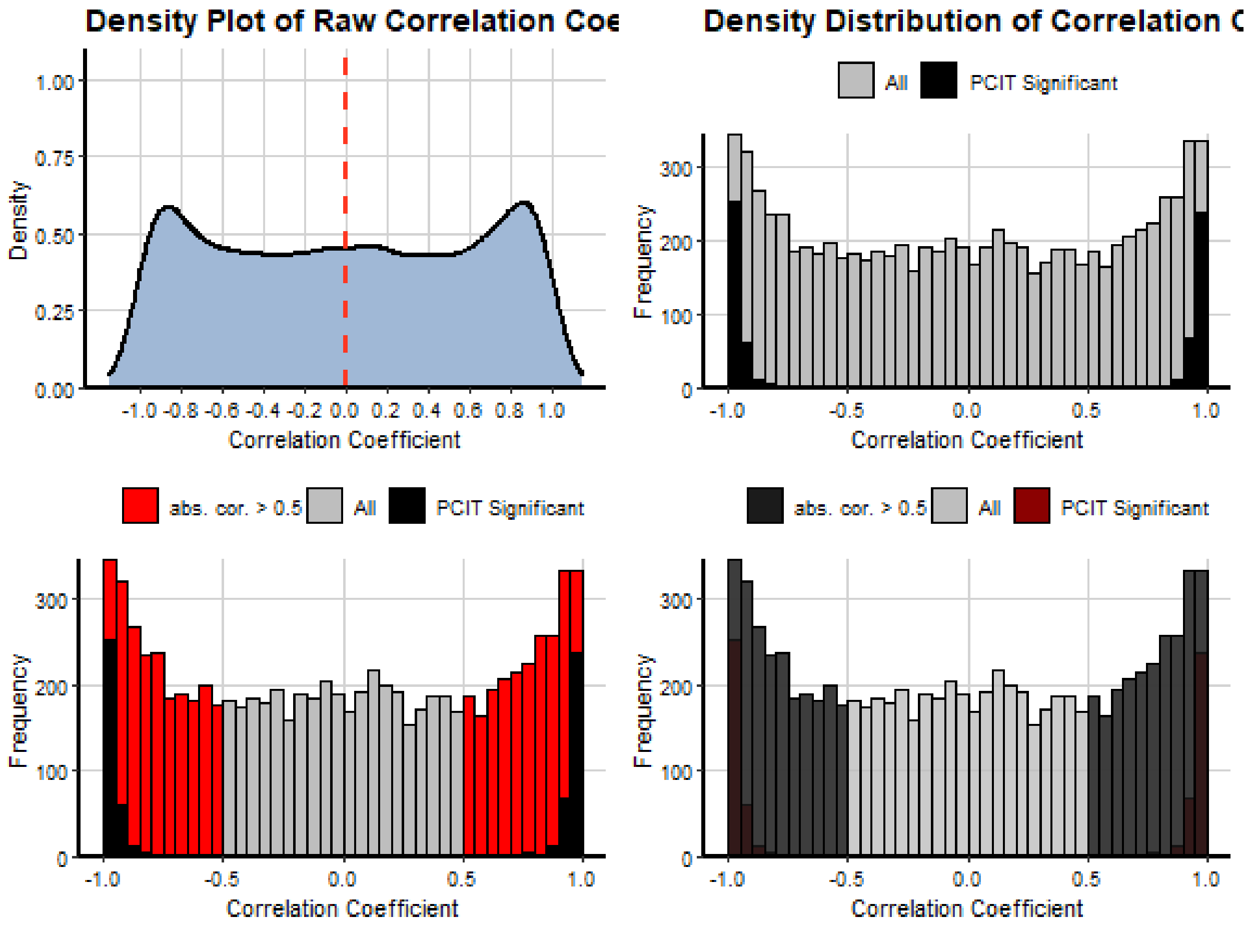


#### RIF

To perform the RIF analysis we will need the count data, an annotation table and a list with the Transcript Factors of specific organism (*Homo sapiens* in this case) and follow the following steps in order to get the output (dataframe with the average expression, RIF1 and RIF2 metrics for each TF):

```r
# Loading packages
library(CeTF)
library(airway)
library(kableExtra)
library(knitr)

# Loading airway data
data("airway")

# Creating a variable with annotation data
anno <- as.data.frame(colData(airway))
```

```r
anno <- anno[order(anno$dex, decreasing = TRUE), ]
anno <- data.frame(cond = anno$dex,
                   row.names = rownames(anno))

# Creating a variable with count data
counts <- assay(airway)

# Sorting count data samples by conditions (untrt and trt)
counts <- counts[, rownames(anno)]
colnames(counts) <- paste0(colnames(counts), c(rep("_untrt", 4), rep("_trt", 4)))

# Differential Expression analysis to use only informative genes
DEGenes <- expDiff(exp = counts,
                   anno = anno,
                   conditions = c('untrt', 'trt'),
                   lfc = 2,
                   padj = 0.05,
                   diffMethod = "Reverter")

# Selecting only DE genes from counts data
counts <- counts[rownames(DEGenes$DE_unique), ]

# Converting count data to TPM
tpm <- apply(counts, 2, function(x) {
            (1e+06 * x)/sum(x)
        })

# Count normalization
Clean_Dat <- normExp(tpm)

# Loading the Transcript Factors (TFs) character
data("TFs")

# Verifying which TFs are in the subsetted normalized data
TFs <- rownames(Clean_Dat)[rownames(Clean_Dat) %in% TFs]

# Selecting the Target genes
Target <- setdiff(rownames(Clean_Dat), TFs)

# Ordering rows of normalized count data
RIF_input <- Clean_Dat[c(Target, TFs), ]

# Performing RIF analysis
RIF_out <- RIF(input = RIF_input,
               nta = length(Target),
               ntf = length(TFs),
               nSamples1 = 4,
               nSamples2 = 4)

# Printing first 10 rows
kable(RIF_out[1:10, ]) %>%
  kable_styling(bootstrap_options = "striped", full_width = FALSE)
```

| TF | avgexpr | RIF1 | RIF2 |
|---|---|---|---|
| ENSG00000008441 | 10.380590 | -0.1386859 | -0.6770894 |
| ENSG00000102804 | 9.264843 | 0.0147001 | 0.1789910 |
| ENSG00000103888 | 13.591598 | 0.0985383 | -0.3361795 |
| ENSG00000103994 | 10.385093 | -1.2748364 | 0.8113721 |
| ENSG00000112837 | 8.143177 | -0.2246323 | -1.7903370 |
| ENSG00000116016 | 11.376028 | -1.2315044 | 0.0216295 |
| ENSG00000116132 | 9.388008 | 1.4038890 | 0.8823288 |
| ENSG00000118689 | 9.189257 | 0.9971123 | 0.3981473 |
| ENSG00000123562 | 9.207009 | 0.3381318 | 0.4609342 |
| ENSG00000157514 | 7.215664 | -1.5628941 | -0.3290858 |

#### Whole analysis of Regulatory Impact Factors (RIF) and Partial Correlation and Information Theory analysis (PCIT)

Finally, it is possible to run the entire analysis all at once. The output will be a CeTF object with all results generated between the different steps. To access the CeTF object is recommended to use the accessors from **CeTF** class:

```
# Loading packages
library(airway)
library(CeTF)

# Loading airway data
data("airway")

# Creating a variable with annotation data
anno <- as.data.frame(colData(airway))

# Creating a variable with count data
counts <- assay(airway)

# Sorting count data samples by conditions (untrt and trt)
counts <- counts[, order(anno$dex, decreasing = TRUE)]

# Loading the Transcript Factors (TFs) character
data("TFs")

# Performing the complete analysis
out <- runAnalysis(mat = counts,
                   conditions=c("untrt", "trt"),
                   lfc = 5,
                   padj = 0.05,
                   TFs = TFs,
                   nSamples1 = 4,
                   nSamples2= 4,
                   tolType = "mean",
                   diffMethod = "Reverter",
                   data.type = "counts")
```

#### Getting some graphical outputs

Based on CeTF class object resulted from **runAnalysis** function is possible to get a plot for Differentially Expressed (DE) genes that shows the relationship between log(baseMean) and Difference of Expression or log2FoldChange, enabling the visualization of the distribution of DE genes and TF in both conditions. These two first plot provides a initial graphical visualization of results:

```
# Using the runAnalysis output (CeTF class object)
SmearPlot(object = out,
          diffMethod = 'Reverter',
          lfc = 1.5,
          conditions = c('untrt', 'trt'),
          type = "DE")
```

```
## Warning: Use of `tab[["baseMean"]]` is discouraged. Use `.data[["baseMean"]]`
## instead.

## Warning: Use of `tab[[var1]]` is discouraged. Use `.data[[var1]]` instead.

## Warning: Use of `tab[["Type"]]` is discouraged. Use `.data[["Type"]]` instead.
```

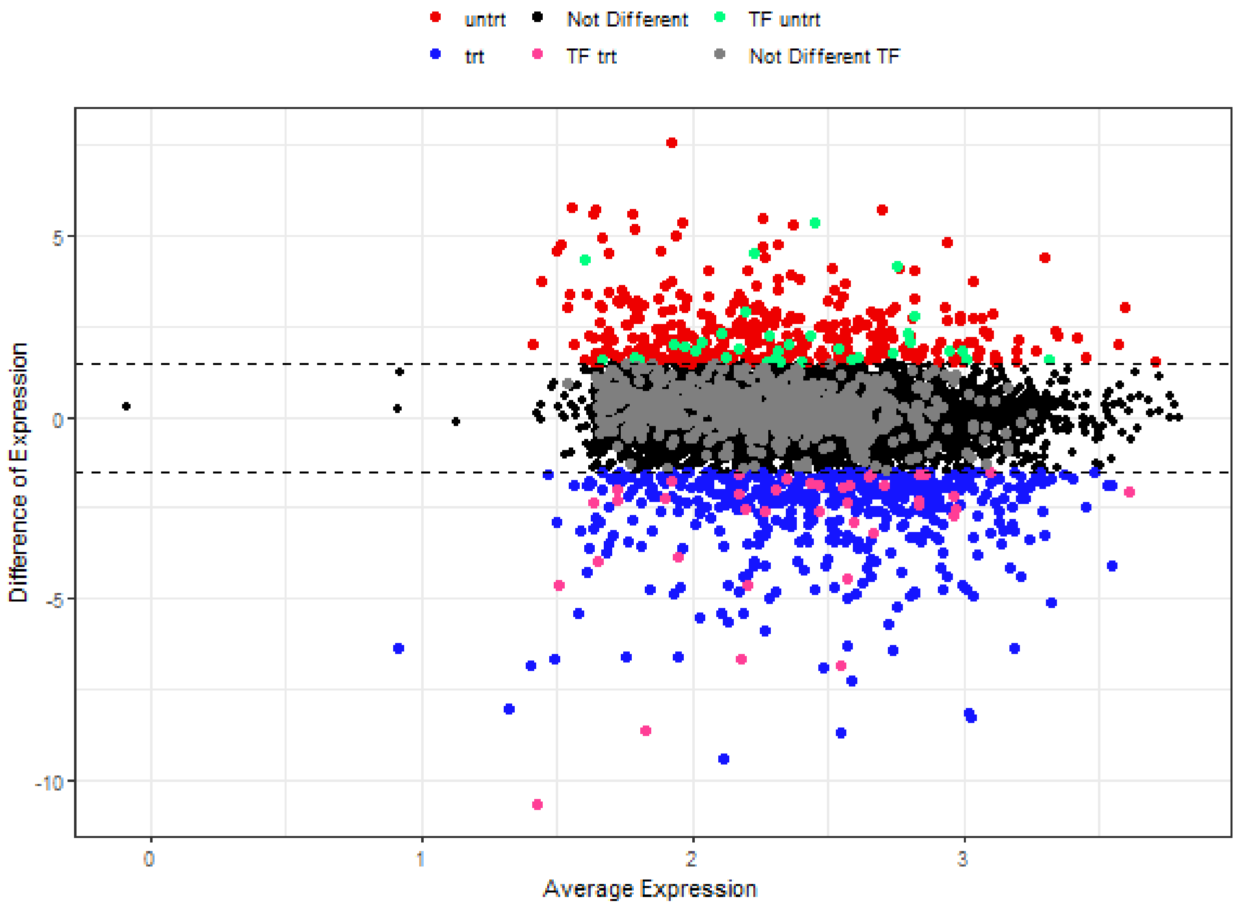


Then is possible to generate the same previous plot but now for a single TF. So, if you have a specific TF that you want to visualize, this is the recommended plot:

```
# Using the runAnalysis output (CeTF class object)
SmearPlot(object = out,
          diffMethod = 'Reverter',
          lfc = 1.5,
          conditions = c('untrt', 'trt'),
          TF = 'ENSG00000185917',
```

```
        label = TRUE,
        type = "TF")
```

```
## Warning: Use of `tab[["baseMean"]]` is discouraged. Use `.data[["baseMean"]]`
## instead.

## Warning: Use of `tab[[var1]]` is discouraged. Use `.data[[var1]]` instead.

## Warning: Use of `tab[["Type"]]` is discouraged. Use `.data[["Type"]]` instead.

## Warning: Use of `dtsub[["baseMean"]]` is discouraged. Use `.data[["baseMean"]]`
## instead.

## Warning: Use of `dtsub[[var1]]` is discouraged. Use `.data[[var1]]` instead.

## Warning: Use of `dtsub[["Type"]]` is discouraged. Use `.data[["Type"]]` instead.

## Warning: Use of `dtsub[["genes"]]` is discouraged. Use `.data[["genes"]]`
## instead.
```

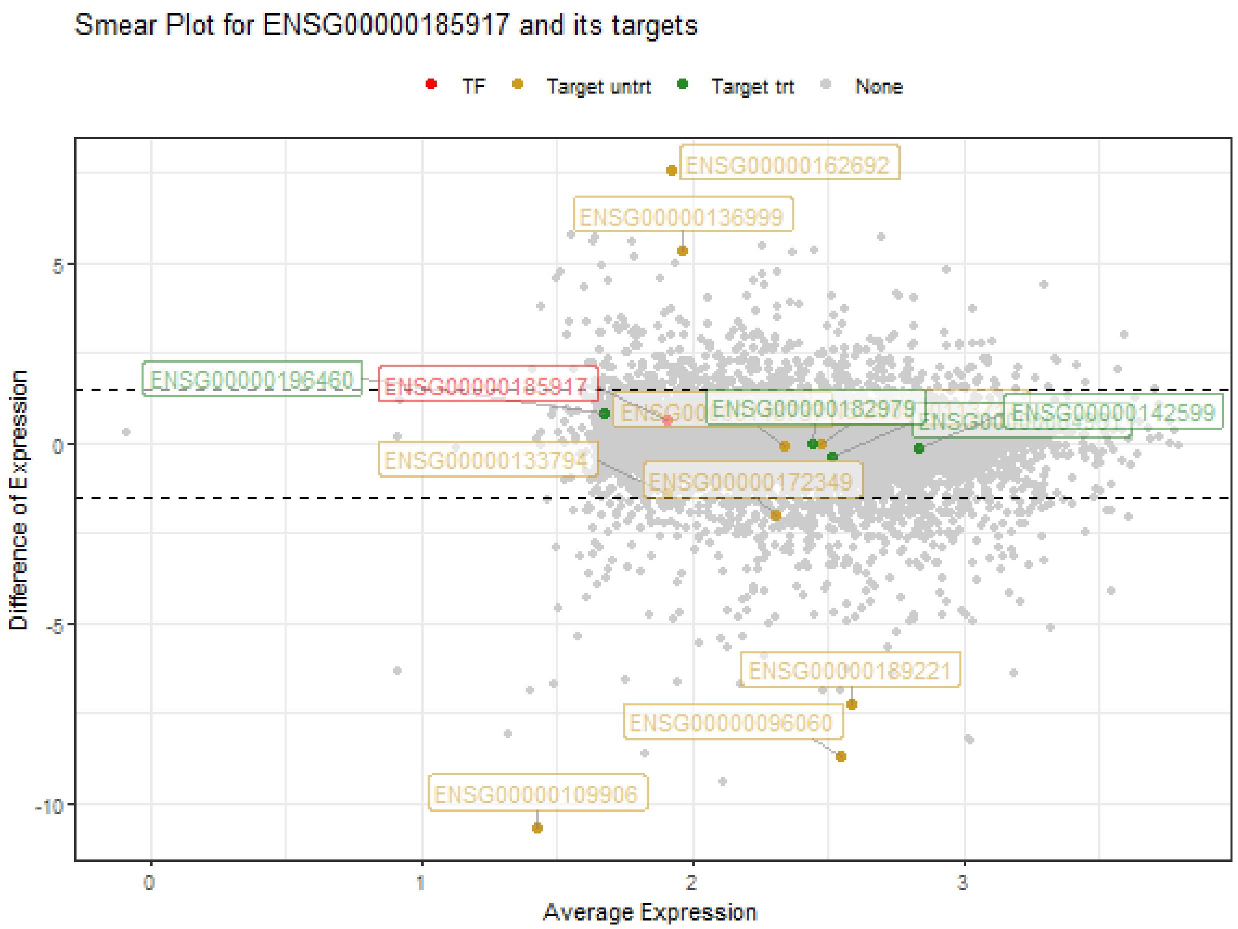


The next output is a network for both conditions resulted from *runAnalysis*:

```
# Using the runAnalysis output (CeTF class object)
netConditionsPlot(out)
```

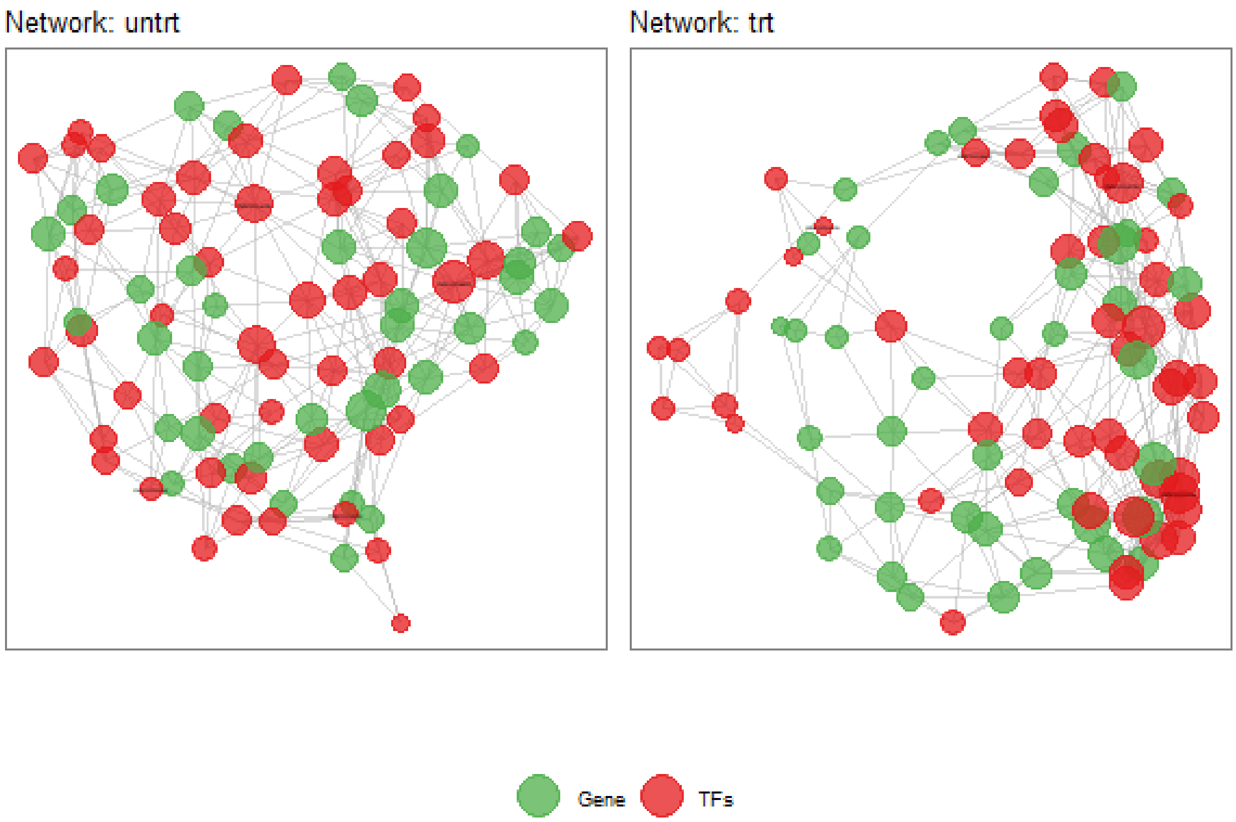


Then we will associate the genes in both conditions with Gene Ontology (GO) and other databases. *getGroupGO* function will be performed where a set of genes are counted within all possible ontologies, without statistical power. This function is capable of perform groupGO for any organism as long as it's annotation package exists (i.e. org.Hs.eg.db, org.Bt.eg.db, org.Rn.eg.db, etc). In this example we will perform only for first condition:

```
# Loading Homo sapiens annotation package
library(org.Hs.eg.db)

# Accessing the network for condition 1
genes <- unique(c(as.character(NetworkData(out, "network1")[, "gene1"]),
                  as.character(NetworkData(out, "network1")[, "gene2"])))

# Performing getGroupGO analysis
cond1 <- getGroupGO(genes = genes,
                    ont = "BP",
                    keyType = "ENSEMBL",
                    annoPkg = org.Hs.eg.db,
                    level = 3)
```

Alternatively, it is possible to perform the enrichment analysis with a statistical power. In this case, there are many databases options to perform this analysis (i.e. GO, KEGG, REACTOME, etc). The *getEnrich* function will perform the enrichment analysis and returns which pathways are enriched in a given set of genes. This analysis requires a reference list of genes with all genes that will be considered to enrich. In this package there is a function, *refGenes* that has these references genes for ENSEMBL and SYMBOL nomenclatures for 5 organisms: Human (**Homo sapiens**), Mouse (**Mus musculus**), Zebrafish (**Danio rerio**), Cow (**Bos taurus**) and Rat (**Rattus norvegicus**). If the user wants to use a different reference set, simply inputs a character vector with the genes.

```
# Accessing the network for condition 1
genes <- unique(c(as.character(NetworkData(out, "network1")[, "gene1"]),
                  as.character(NetworkData(out, "network1")[, "gene2"])))

# Performing getEnrich analysis
enrich <- getEnrich(organism='hsapiens', database='geneontology_Biological_Process',
                   genes=genes, GeneType='ensembl_gene_id',
                   refGene=refGenes$Homo_sapiens$ENSEMBL,
                   fdrMethod = "BH", fdrThr = 0.05, minNum = 5, maxNum = 500)
```

In the next steps we will use the *getGroupGO* results, but it is worth mentioning that you can use the results of the *getEnrich* function. To run the following steps with the *getEnrich* results is recommended to change the *lfc* parameter in aboce *runAnalysis* function by 3. That way there will be more pathways to enrich. Remembering that this will require more processing time.

After the association of GOs and genes, we are able to plot the network of previously results. It's possible to choose which variables ("pathways", "TFs" or "genes") we want to group. All these variables are inputted by the user, i.e. the user can choose the pathways, TFs or genes returned from analysis from the runAnalysis function and CeTF class object, or can also choose by specific pathways, TFs or genes of interest. The table used to plot the networks are stored in a list resulted from netGOTFPlot function. To get access is needed to use the *object$tab*. The list will be splitted by groupBy argument choice:

```
# Subsetting only the first 12 Ontologies with more counts
t1 <- head(cond1$results, 12)

# Subsetting the network for the conditions to make available only the 12 nodes subsetted
t2 <- subset(cond1$netGO, cond1$netGO$gene1 %in% as.character(t1[, "ID"]))

# generating the GO plot grouping by pathways
pt <- netGOTFPlot(netCond = NetworkData(out, "network1"),
                  resultsGO = t1,
                  netGO = t2,
                  anno = NetworkData(out, "annotation"),
                  groupBy = 'pathways',
                  type = 'GO')
pt$plot
```

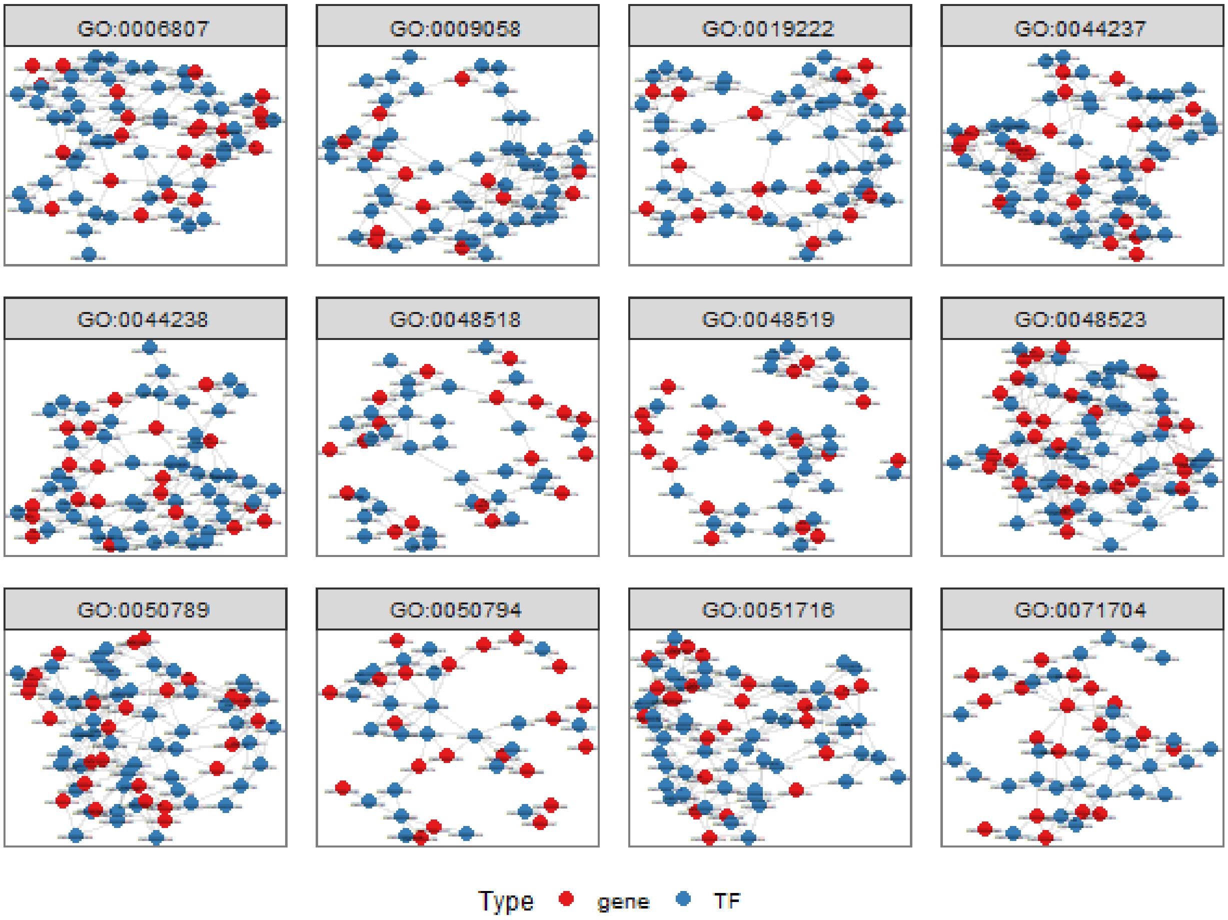


```
head(pt$tab$`GO:0006807`)
```

```
##                 gene1    pathway           gene2 class
## 1   ENSG00000004799 GO:0006807 ENSG00000114861  gene
## 2   ENSG00000004799 GO:0006807 ENSG00000119138  gene
## 3   ENSG00000004799 GO:0006807 ENSG00000120093  gene
## 4   ENSG00000004799 GO:0006807 ENSG00000184271  gene
## 52  ENSG00000013293 GO:0019222 ENSG00000113916  gene
## 109 ENSG00000030419 GO:0009058 ENSG00000170448    TF
```

```
# generating the GO plot grouping by TFs
TFs <- NetworkData(out, "keytfs")$TF[1:4]
pt <- netGOTFPlot(netCond = NetworkData(out, "network1"),
                  resultsGO = t1,
                  netGO = t2,
                  anno = NetworkData(out, "annotation"),
                  groupBy = 'TFs',
                  TFs = TFs,
                  type = 'GO')
pt$plot
```

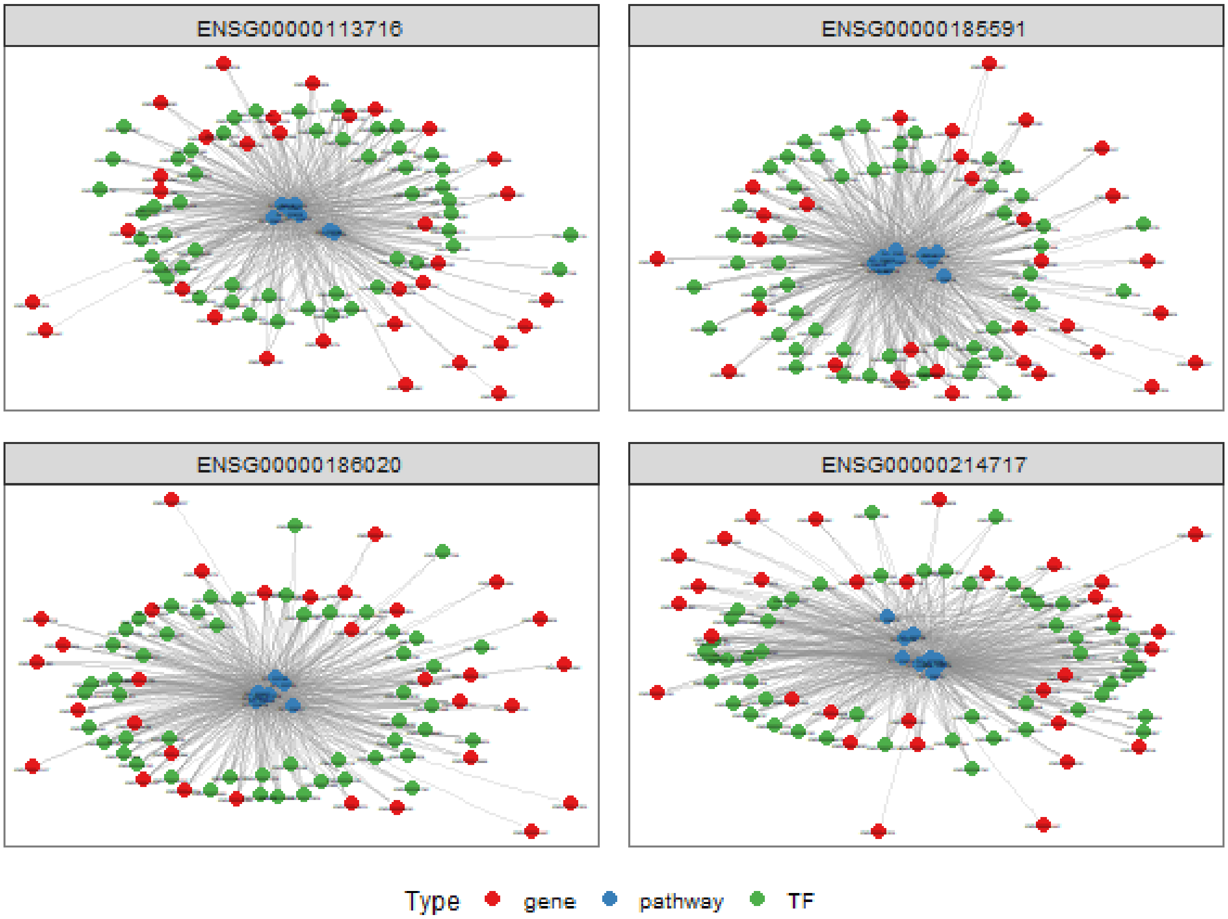


```
head(pt$tab$ENSG00000011465)
```

```
## NULL
```

```
# generating the GO plot grouping by specifics genes
genes <- c('ENSG00000011465', 'ENSG00000026025', 'ENSG00000075624', 'ENSG00000077942')
pt <- netGOTFPlot(netCond = NetworkData(out, "network1"),
                  resultsGO = t1,
                  netGO = t2,
                  anno = NetworkData(out, "annotation"),
                  groupBy = 'genes',
                  genes = genes,
                  type = 'GO')
pt$plot
```

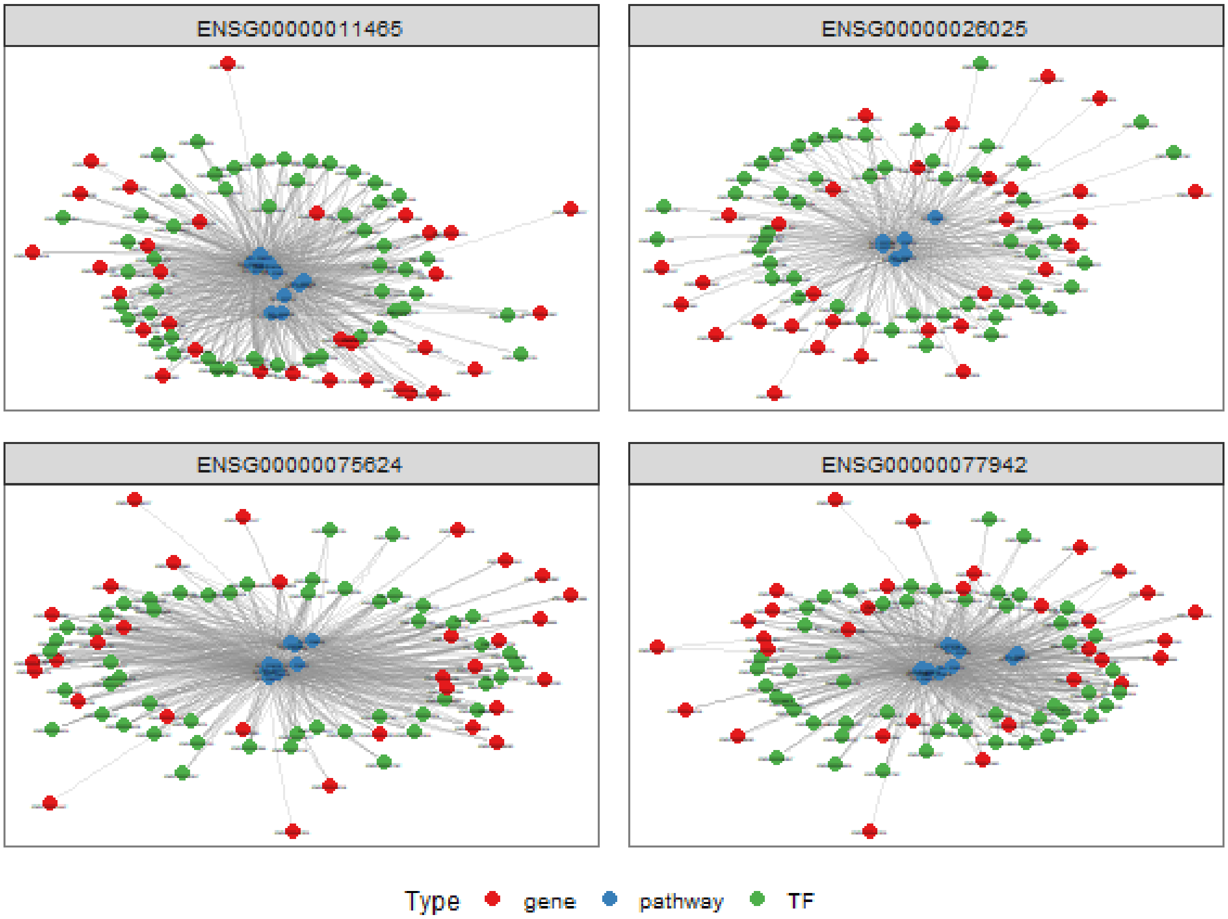


```r
head(pt$tab$ENSG00000011465)
```

```
##        gene1           genes           gene2   class
## 1 GO:0050789 ENSG00000011465 ENSG00000004799 pathway
## 2 GO:0050789 ENSG00000011465 ENSG00000013293 pathway
## 3 GO:0050789 ENSG00000011465 ENSG00000030419 pathway
## 4 GO:0050789 ENSG00000011465 ENSG00000053254 pathway
## 5 GO:0050789 ENSG00000011465 ENSG00000064961 pathway
## 6 GO:0050789 ENSG00000011465 ENSG00000067082 pathway
```

Finally, we will merge the results from the network of condition 1 (Gene-Gene and TF-Gene interaction) with groupGO results (GO-Gene or GO-TF interaction). The final results has the objective to integrate all these data in order to results in a complete view of the data:

```r
pt <- netGOTFPlot(netCond = NetworkData(out, "network1"),
                  netGO = t2,
                  keyTFs = NetworkData(out, "keytfs"),
                  type = 'Integrated')

pt$plot
```

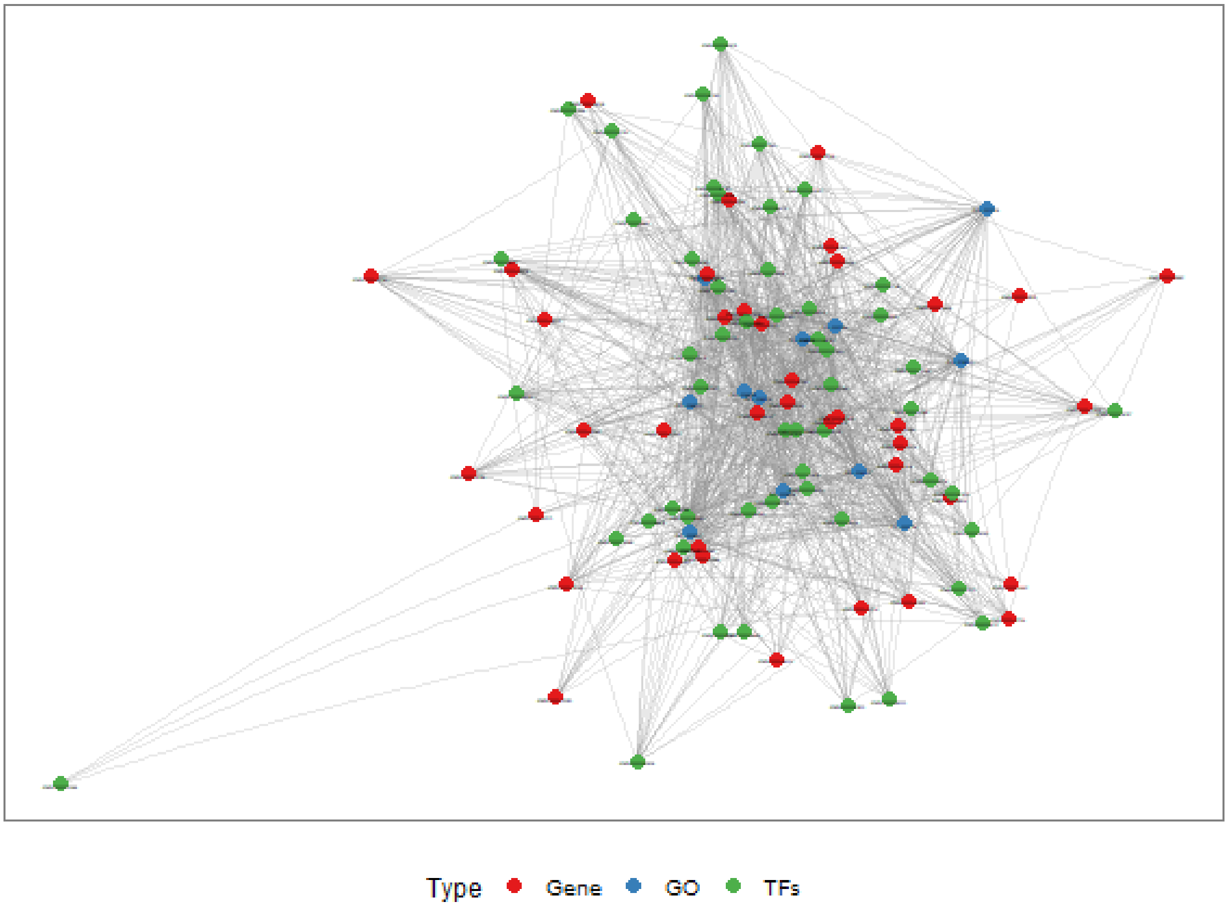


```r
head(pt$tab)
```

```
##                gene1           gene2
## 22 ENSG00000004799 ENSG00000113761
## 24 ENSG00000004799 ENSG00000114861
## 26 ENSG00000004799 ENSG00000119138
## 27 ENSG00000004799 ENSG00000120093
## 47 ENSG00000004799 ENSG00000143127
## 77 ENSG00000004799 ENSG00000184271
```

#### Using accessors to access results

Is possible to use the accessors from CeTF class object to access the outputs generated by the *runAnalysis* function. These outputs can be used as input in **Cytoscape** (Shannon et al. 2003). The accessors are:

- **getData**: returns the raw, tpm and normalized data;
- **getDE**: returns the DE genes and TFs;
- **InputData** returns the input matrices used to perform RIF and PCIT in *runAnalysis* function;
- **OutputData** returns the output matrices and lists that output from RIF and PCIT in *runAnalysis* function;
- **NetworkData** returns the network of interactions between gene and key TFs for both conditions, the keyTFs and the annotation for each gene and TF.

All accessors can be further explored by looking in more detail at the documentation.

### Additional Features

This package has some additional features to plot the results. Let's use the gene set from *airway* data previously used. The features are:

#### Network diffusion analysis

Network propagation is an important and widely used algorithm in systems biology, with applications in protein function prediction, disease gene prioritization, and patient stratification. Propagation provides a robust estimate of network distance between sets of nodes. Network propagation uses the network of interactions to find new genes that are most relevant to a set of well understood genes. To perform this analysis is necessary to install the latest Cytoscape software version and know the path that will be installed the software. After running the diffusion analysis is possible to perform the enrichment for the subnetwork resulted. Finally, ih this vignette we'll not perform the diffusion analysis because this requires Cytoscape to be installed.

```
result <- diffusion(object = out,
                    cond = "network1",
                    genes = c("ENSG00000185591", "ENSG00000179094"),
                    cyPath = "C:/Program Files/Cytoscape_v3.7.2/Cytoscape.exe",
                    name = "top_diffusion",
                    label = T)
```

#### Circos plot

This plot makes it possible to visualize the targets of specific TFs or genes. The main idea is to identify which chromosomes are linked to the targets of a given gene or TF to infer whether there are cis (same chromosome) or trans (different chromosomes) links between them. The black links are between different chromosomes while the red links are between the same chromosome.

```
# Loading the CeTF object class demo
data("CeTFdemo")

CircosTargets(object = CeTFdemo,
              file = "/path/to/gtf/file/specie.gtf",
              nomenclature = "ENSEMBL",
              selection = "ENSG00000185591",
              cond = "condition1")
```

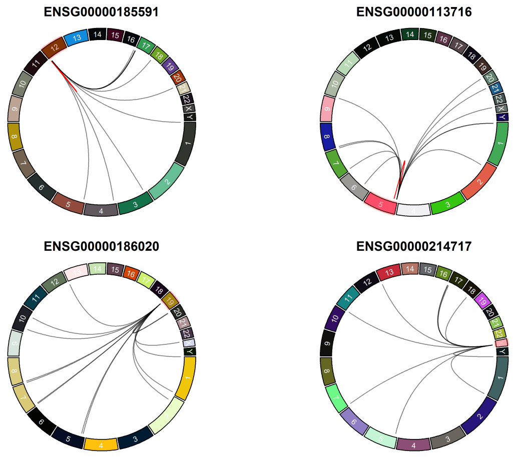

##### RIF relationships plots

To visualize the relationship between RIF results (RIF1 and RIF2) is possible to generate the following plot:

```
RIFPlot(object = out,
        color  = "darkblue",
        type   = "RIF")
```

```
## Warning: Use of `tmp[["RIF1"]]` is discouraged. Use `.data[["RIF1"]]` instead.
## Warning: Use of `tmp[["RIF2"]]` is discouraged. Use `.data[["RIF2"]]` instead.
```

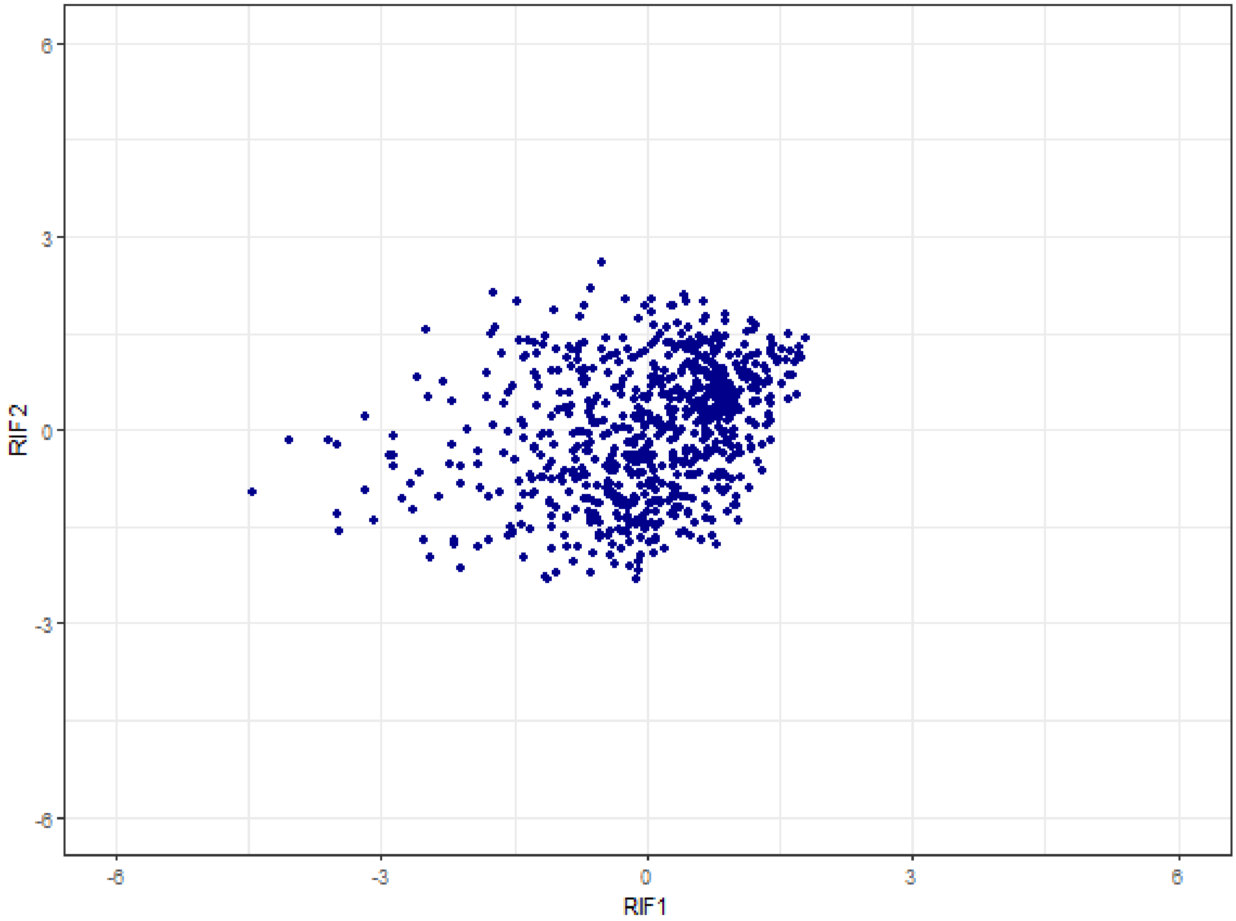


Then to get the relationship between RIF1/RIF2 and differential expressed genes:

```
RIFPlot(object = out,
        color  = "darkblue",
        type   = "DE")
```

```
## Warning: Use of `tb[["RIF1"]]` is discouraged. Use `.data[["RIF1"]]` instead.
## Warning: Use of `tb[["DE"]]` is discouraged. Use `.data[["DE"]]` instead.
## Warning: Use of `tb[["RIF2"]]` is discouraged. Use `.data[["RIF2"]]` instead.
## Warning: Use of `tb[["DE"]]` is discouraged. Use `.data[["DE"]]` instead.
```

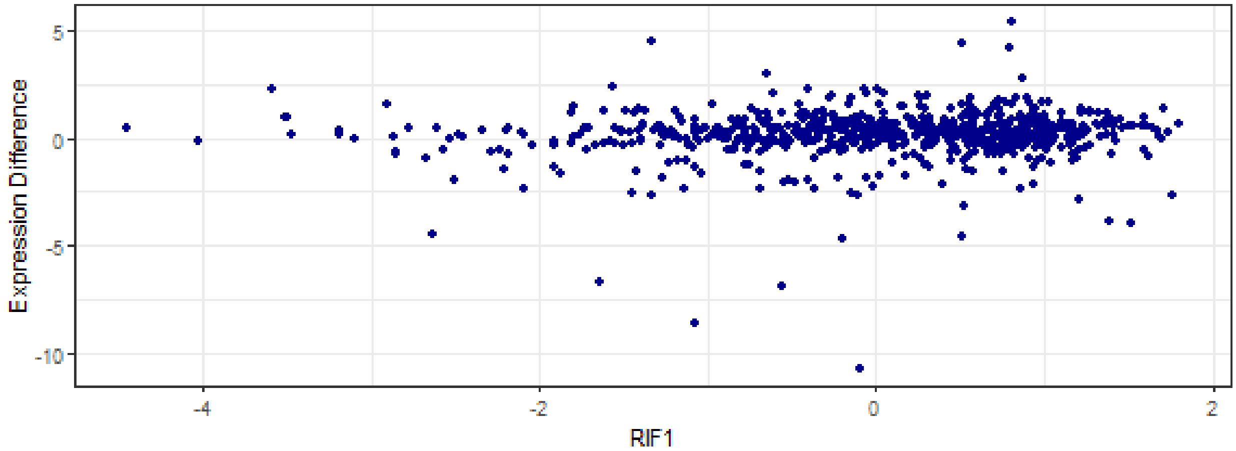


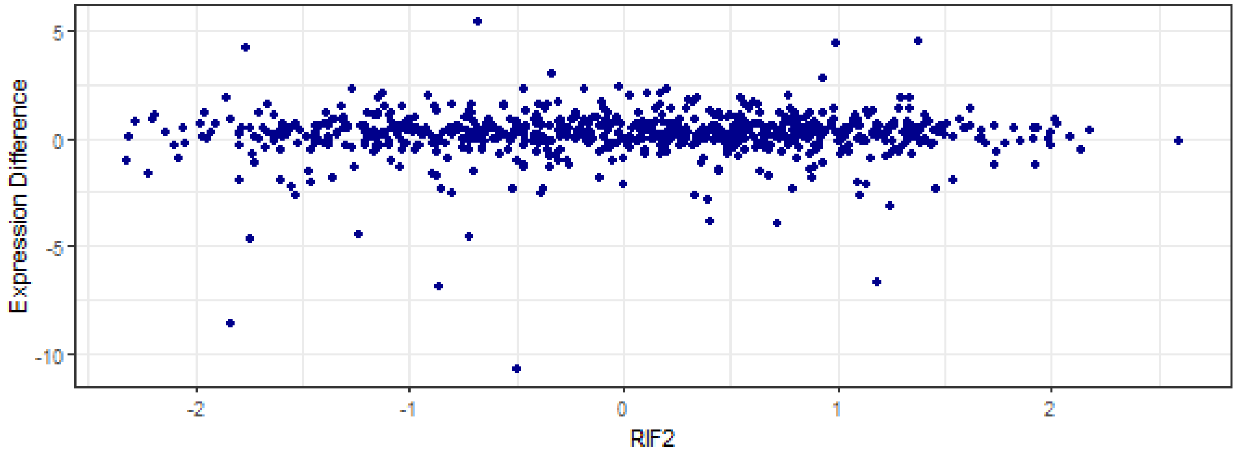


##### Enrichment plots

We provide some option to plot the results from **getEnrich** function. First of all we need to perform the enrichment analysis for a set of genes:

```
# Selecting the genes related to the network for condition 1
genes <- unique(c(as.character(NetworkData(out, "network1")[, "gene1"]),
                  as.character(NetworkData(out, "network1")[, "gene2"])))

# Performing enrichment analysis
cond1 <- getEnrich(organism='hsapiens', database='geneontology_Biological_Process',
                   genes=genes, GeneType='ensembl_gene_id',
                   refGene=refGenes$Homo_sapiens$ENSEMBL,
                   fdrMethod = "BH", fdrThr = 0.05, minNum = 5, maxNum = 500)
```

```
## Loading the functional categories...
## Loading the ID list...
## Loading the reference list...
## Performing the enrichment analysis...
```

After performing the enrichment analysis we have 4 options for viewing this data:

##### Heatmap-like functional classification

```r
heatPlot(res = cond1$results,
         diff = getDE(out, "all"))
```

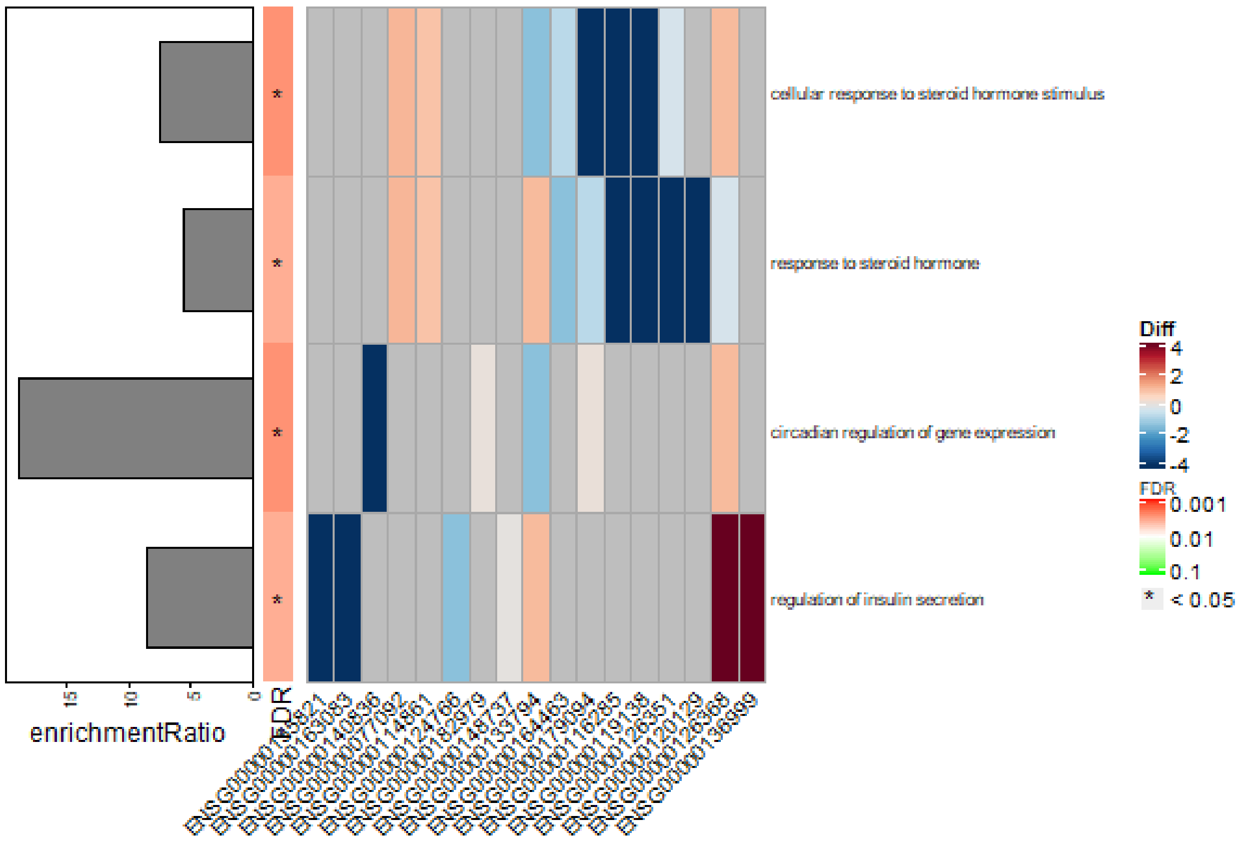


##### Circle Barplot

```r
enrichPlot(res = cond1$results,
           type = "circle")
```

```
## Warning: Use of `res[["enrichmentRatio"]]` is discouraged. Use
## `.data[["enrichmentRatio"]]` instead.

## Warning: Use of `res[["FDR"]]` is discouraged. Use `.data[["FDR"]]` instead.

## Warning: Use of `res[["description"]]` is discouraged. Use
## `.data[["description"]]` instead.

## Warning: Use of `res[["enrichmentRatio"]]` is discouraged. Use
## `.data[["enrichmentRatio"]]` instead.
```

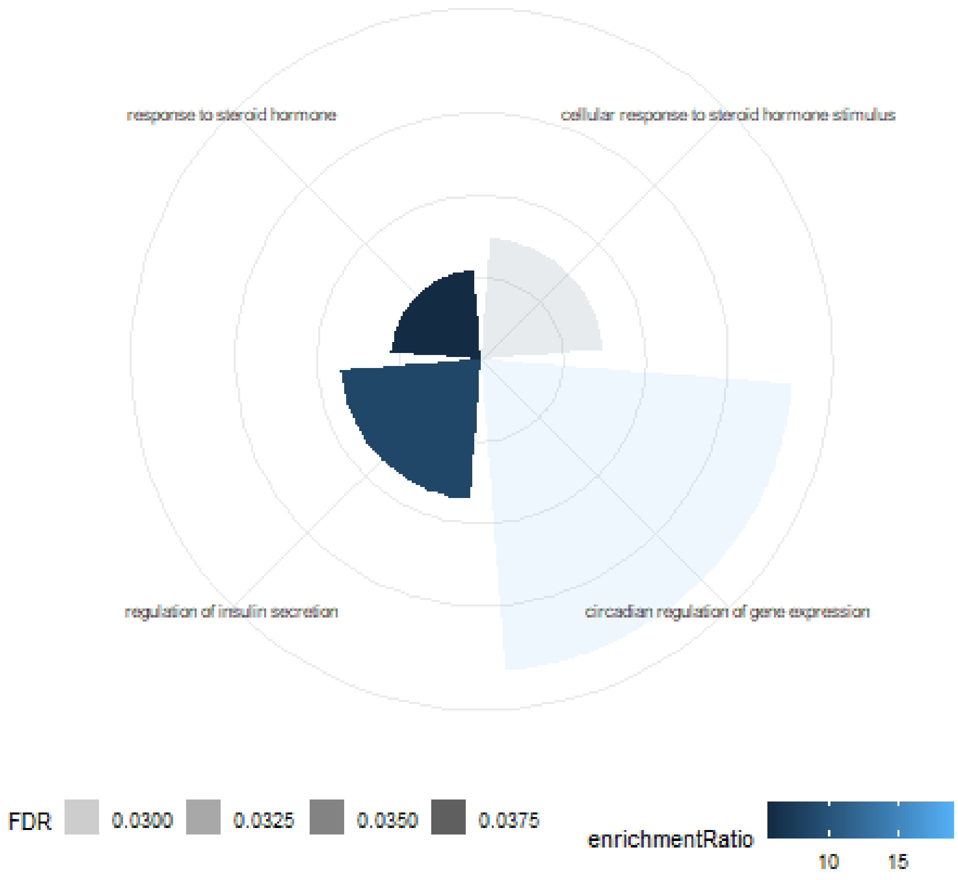


##### Barplot

```
enrichPlot(res = cond1$results,
           type = "bar")
```

```
## Warning: Use of `res[["FDR"]]` is discouraged. Use `.data[["FDR"]]` instead.
```

```
## Warning: Use of `res[["description"]]` is discouraged. Use
## `.data[["description"]]` instead.
```

```
## Warning: Use of `res[["enrichmentRatio"]]` is discouraged. Use
## `.data[["enrichmentRatio"]]` instead.
```

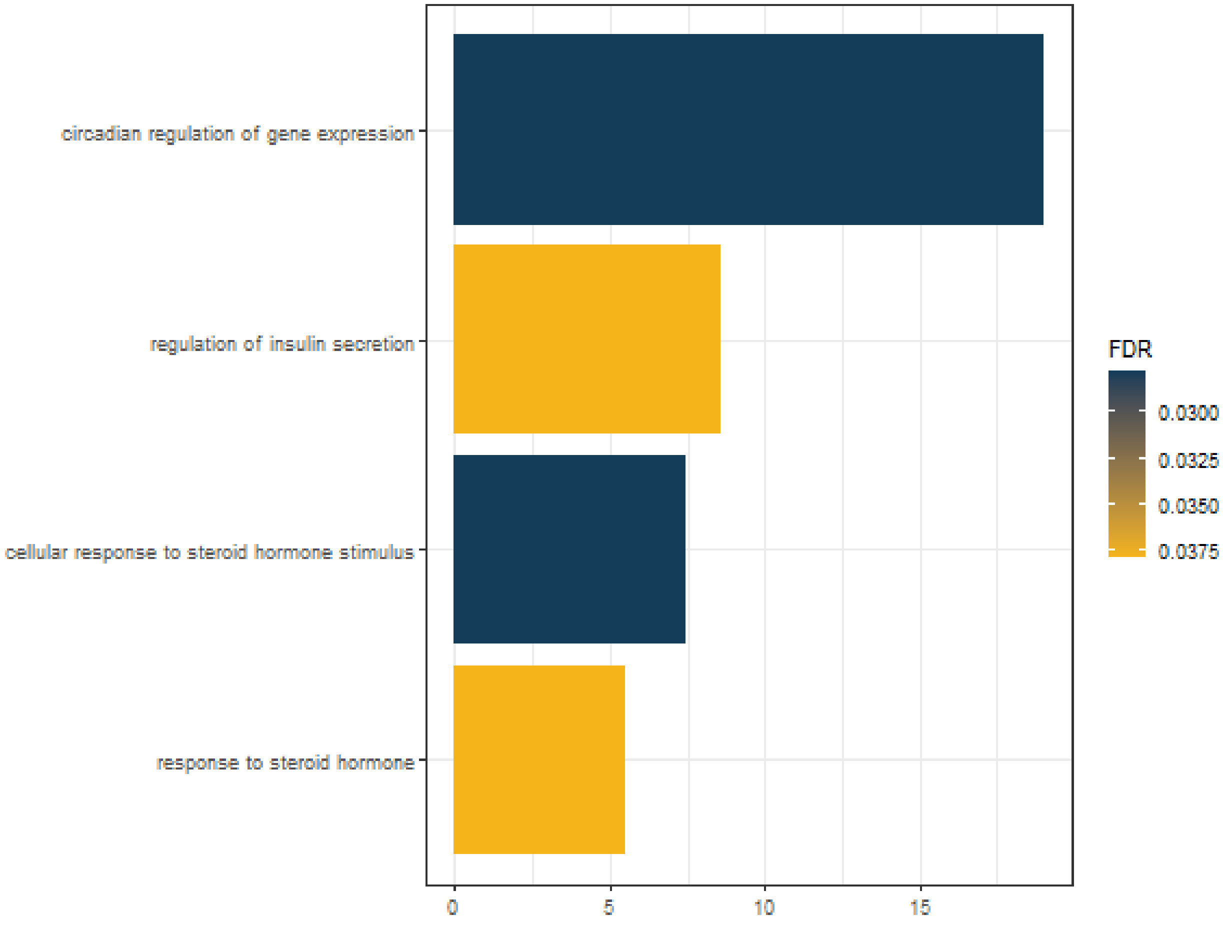


**Dotplot**

```
enrichPlot(res = cond1$results,
           type = "dot")
```

```
## Warning: Use of `res[["description"]]` is discouraged. Use
## `.data[["description"]]` instead.

## Warning: Use of `res[["enrichmentRatio"]]` is discouraged. Use
## `.data[["enrichmentRatio"]]` instead.

## Warning: Use of `res[["overlap"]]` is discouraged. Use `.data[["overlap"]]`
## instead.

## Warning: Use of `res[["FDR"]]` is discouraged. Use `.data[["FDR"]]` instead.
```

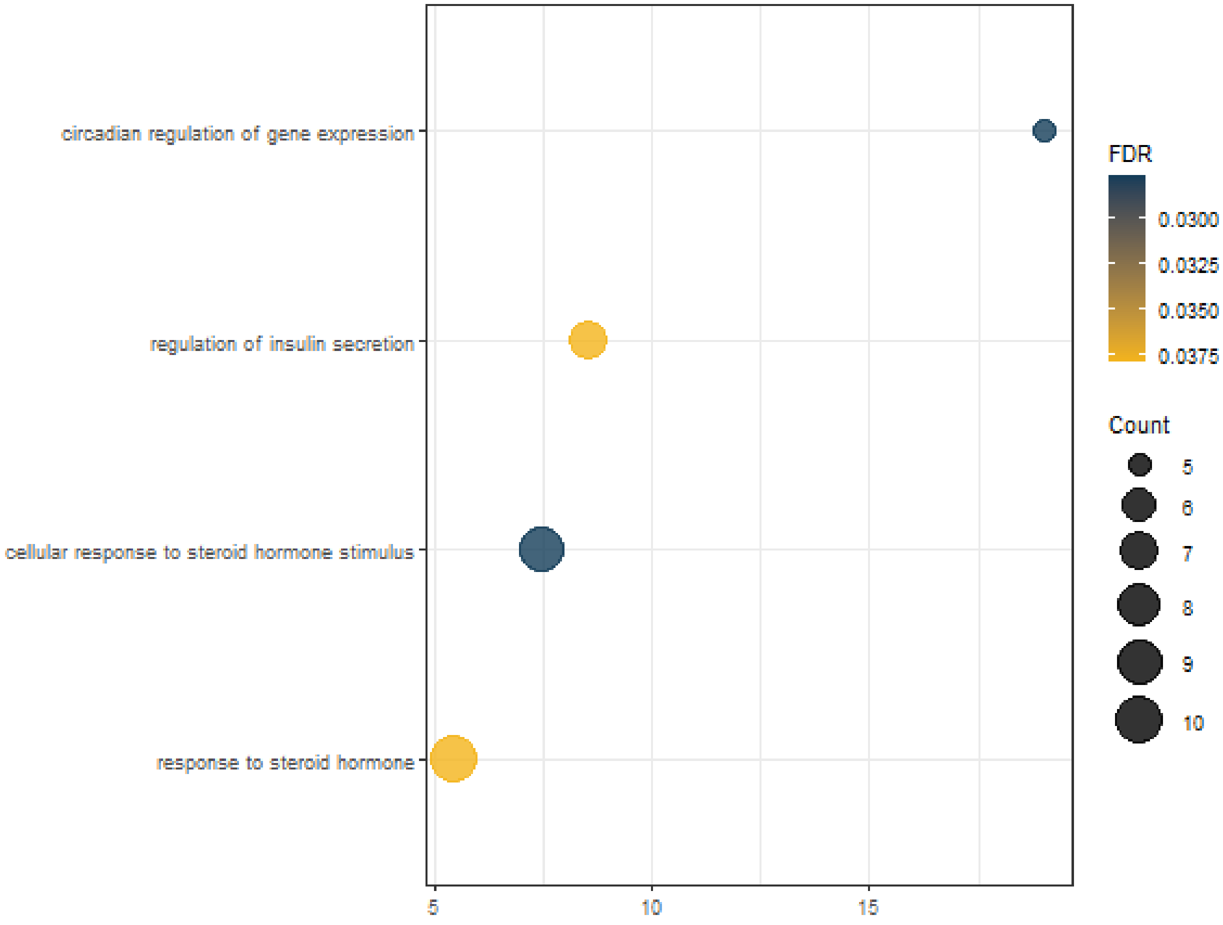


#### Session info

```
sessionInfo()
```

```
## R Under development (unstable) (2020-01-07 r77637)
## Platform: x86_64-w64-mingw32/x64 (64-bit)
## Running under: Windows 10 x64 (build 18363)
##
## Matrix products: default
##
## locale:
## [1] LC_COLLATE=Portuguese_Brazil.1252  LC_CTYPE=Portuguese_Brazil.1252
## [3] LC_MONETARY=Portuguese_Brazil.1252 LC_NUMERIC=C
## [5] LC_TIME=Portuguese_Brazil.1252
##
## attached base packages:
## [1] parallel  stats4    stats     graphics  grDevices utils     datasets
## [8] methods   base
##
## other attached packages:
##  [1] org.Hs.eg.db_3.10.0         AnnotationDbi_1.49.1
##  [3] knitr_1.28                  kableExtra_1.1.0
##  [5] airway_1.7.0                SummarizedExperiment_1.17.3
```

```
##  [7] DelayedArray_0.13.4         BiocParallel_1.21.2
##  [9] matrixStats_0.56.0          Biobase_2.47.3
## [11] GenomicRanges_1.39.2        GenomeInfoDb_1.23.13
## [13] IRanges_2.21.5              S4Vectors_0.25.13
## [15] BiocGenerics_0.33.2         CeTF_0.99.16
##
## loaded via a namespace (and not attached):
##   [1] circlize_0.4.8                    fastmatch_1.1-0
##   [3] systemfonts_0.1.1                 plyr_1.8.6
##   [5] igraph_1.2.4.2                    RCy3_2.7.4
##   [7] splines_4.0.0                     ggnetwork_0.5.8
##   [9] ggplot2_3.3.0                     urltools_1.7.3
##  [11] digest_0.6.25                     foreach_1.4.8
##  [13] htmltools_0.4.0                   GOSemSim_2.13.1
##  [15] viridis_0.5.1                     GO.db_3.10.0
##  [17] magrittr_1.5                      memoise_1.1.0
##  [19] cluster_2.1.0                     doParallel_1.0.15
##  [21] GenomicTools_0.2.9.7              sna_2.5
##  [23] ComplexHeatmap_2.3.2              readr_1.3.1
##  [25] annotate_1.65.1                   graphlayouts_0.6.0
##  [27] R.utils_2.9.2                     svglite_1.2.3
##  [29] prettyunits_1.1.1                 enrichplot_1.7.3
##  [31] colorspace_1.4-1                  rvest_0.3.5
##  [33] blob_1.2.1                        ggrepel_0.8.2
##  [35] xfun_0.12                         dplyr_0.8.5
##  [37] crayon_1.3.4                      RCurl_1.98-1.1
##  [39] jsonlite_1.6.1                    scatterpie_0.1.4
##  [41] graph_1.65.2                      genefilter_1.69.0
##  [43] survival_3.1-8                    iterators_1.0.12
##  [45] glue_1.3.2                        polyclip_1.10-0
##  [47] gtable_0.3.0                      zlibbioc_1.33.1
##  [49] XVector_0.27.1                    webshot_0.5.2
##  [51] GetoptLong_0.1.8                  shape_1.4.4
##  [53] apcluster_1.4.8                   scales_1.1.0
##  [55] DOSE_3.13.2                       mvtnorm_1.1-0
##  [57] DBI_1.1.0                         GGally_1.4.0
##  [59] rngtools_1.5                      Rcpp_1.0.4
##  [61] GenomicTools.fileHandler_0.1.5.9 progress_1.2.2
##  [63] viridisLite_0.3.0                 xtable_1.8-4
##  [65] clue_0.3-57                       gridGraphics_0.5-0
##  [67] europepmc_0.3                     bit_1.1-15.2
##  [69] httr_1.4.1                        fgsea_1.13.4
##  [71] RColorBrewer_1.1-2                pkgconfig_2.0.3
##  [73] reshape_0.8.8                     XML_3.99-0.3
##  [75] R.methodsS3_1.8.0                 farver_2.0.3
##  [77] locfit_1.5-9.1                    RJSONIO_1.3-1.4
##  [79] labeling_0.3                      ggplotify_0.0.5
##  [81] clinfun_1.0.15                    tidyselect_1.0.0
##  [83] rlang_0.4.5                       reshape2_1.4.3
##  [85] munsell_0.5.0                     tools_4.0.0
##  [87] downloader_0.4                    RSQLite_2.2.0
##  [89] statnet.common_4.3.0              ggridges_0.5.2
##  [91] evaluate_0.14                     stringr_1.4.0
##  [93] yaml_2.2.1                        WebGestaltR_0.4.3
```

```
##  [95] bit64_0.9-7                    tidygraph_1.1.2
##  [97] purrr_0.3.3                    gMWT_1.1.1
##  [99] ggraph_2.0.2                   pbapply_1.4-2
## [101] doRNG_1.8.2                    whisker_0.4
## [103] R.oo_1.23.0                    xml2_1.2.5
## [105] DO.db_2.9                      rstudioapi_0.11
## [107] compiler_4.0.0                 curl_4.3
## [109] png_0.1-7                      ggsignif_0.6.0
## [111] tibble_2.1.3                   tweenr_1.0.1
## [113] geneplotter_1.65.0             stringi_1.4.6
## [115] gdtools_0.2.1                  lattice_0.20-38
## [117] Matrix_1.2-18                  vctrs_0.2.4
## [119] pillar_1.4.3                   lifecycle_0.2.0
## [121] BiocManager_1.30.10            triebeard_0.3.0
## [123] GlobalOptions_0.1.1            snpStats_1.37.0
## [125] cowplot_1.0.0                  data.table_1.12.8
## [127] bitops_1.0-6                   qvalue_2.19.0
## [129] R6_2.4.1                       network_1.16.0
## [131] gridExtra_2.3                  codetools_0.2-16
## [133] MASS_7.3-51.5                  assertthat_0.2.1
## [135] DESeq2_1.27.29                 rjson_0.2.20
## [137] GenomeInfoDbData_1.2.2         hms_0.5.3
## [139] clusterProfiler_3.15.4         grid_4.0.0
## [141] coda_0.19-3                    tidyr_1.0.2
## [143] rvcheck_0.1.8                  rmarkdown_2.1
## [145] ggpubr_0.2.5                   ggforce_0.3.1
```

### References


Cover, Thomas M, and Joy A Thomas. 2012. *Elements of Information Theory*. John Wiley & Sons.

Himes, Blanca E, Xiaofeng Jiang, Peter Wagner, Ruoxi Hu, Qiyu Wang, Barbara Klanderman, Reid M Whitaker, et al. 2014. "RNA-Seq Transcriptome Profiling Identifies Crispld2 as a Glucocorticoid Responsive Gene That Modulates Cytokine Function in Airway Smooth Muscle Cells." *PloS One* 9 (6): e99625.

Reverter, Antonio, and Eva KF Chan. 2008. "Combining Partial Correlation and an Information Theory Approach to the Reversed Engineering of Gene Co-Expression Networks." *Bioinformatics* 24 (21): 2491–7.

Reverter, Antonio, Nicholas J Hudson, Shivashankar H Nagaraj, Miguel Pérez-Enciso, and Brian P Dalrymple. 2010. "Regulatory Impact Factors: Unraveling the Transcriptional Regulation of Complex Traits from Expression Data." *Bioinformatics* 26 (7): 896–904.

Shannon, Paul, Andrew Markiel, Owen Ozier, Nitin S Baliga, Jonathan T Wang, Daniel Ramage, Nada Amin, Benno Schwikowski, and Trey Ideker. 2003. "Cytoscape: A Software Environment for Integrated Models of Biomolecular Interaction Networks." *Genome Research* 13 (11): 2498–2504.

Vaquerizas, Juan M, Sarah K Kummerfeld, Sarah A Teichmann, and Nicholas M Luscombe. 2009. "A Census of Human Transcription Factors: Function, Expression and Evolution." *Nature Reviews Genetics* 10 (4): 252.

Wang, Kai, and Hiroki Nishida. 2015. "REGULATOR: A Database of Metazoan Transcription Factors and Maternal Factors for Developmental Studies." *BMC Bioinformatics* 16 (1): 114.
